# Neural processing of bottom up perception of biological motion under attentional load

**DOI:** 10.1101/2023.03.14.532555

**Authors:** Hilal Nizamoglu, Burcu A. Urgen

## Abstract

Considering its importance for one’s survival and social significance, biological motion (BM) perception is assumed to occur automatically. Indeed, Thornton and Vuong (2004) showed that task-irrelevant BM in the periphery interfered with task performance at the fovea. However, the neural underpinnings of this bottom-up processing of BM lacks thorough examination in the field. Under selective attention, BM perception is supported by a network of regions including the occipito-temporal, parietal, and premotor cortices. A retinotopy mapping study on BM showed distinct maps for its processing under and away from selective attention (Saygin & Sereno, 2008). Based on these findings, we investigated how bottom-up perception of BM would be processed in the human brain under attentional load when it was shown away from the focus of attention as a task-irrelevant stimulus. Participants (N=31) underwent an fMRI study in which they performed an attentionally demanding visual detection task at the fovea while intact or scrambled PLDs of BM were shown at the periphery. Our results showed the main effect of attentional load in fronto-parietal regions and the main effect of peripheral stimuli in occipito-temporal cortex. Both univariate activity maps and multivariate pattern analysis results support the attentional load modulation on BM in the occipito-temporal cortex. In conclusion, BM is processed within the motion sensitive regions in the occipito-temporal cortex even when it is away from selective attention, and is modulated by the top-down factor of attentional load.

## 1. Introduction

Humans and non-human animals are considered to have an innate tendency to detect and discriminate the movements of other living things from the movements of non-living things (such as objects) starting from the early ages of life (Bertenthal, Proffitt, & Kramer, 1987; Fox & McDaniel, 1982; Pavlova, Krageloh-Mann, Sokolov, & Birbaumer, 2001; Sifre et al., 2018; Simion, Regolin & Bulf, 2008; Vallortigara et al., 2005). Besides its obvious importance for survival, this skill, also known as the ability to perceive biological motion (BM), constitutes the basis for higher order social skills such as communication and social interaction in humans (Blake & Shiffrar, 2007). By simply observing a person’s BM, one can infer the emotional state (Halovic & Kroos, 2018), gender (Pollick, Kay, Heim, & Stringer, 2005), and age of the person (see Blake & Shiffrar, 2007) as well as their social characteristics such as identity (Yovel & O’Toole, 2016) and personal traits (Thoresen, Vuong & Atkinson, 2012).

According to Thompson and Parasuraman (2012), BM perception requires three main steps of processing: (i) detecting the body features constituting a movement; (ii) forming action representations; and (iii) understanding the intentions conveyed from observed actions. These steps are supported by a network of regions including occipito-temporal cortex (OTC) regions that involve form and motion sensitive areas, including extrastriate body area (EBA) and human middle temporal cortex cluster (hMT+) (Pyles & Grossman, 2013; Downing et al., 2001; Grossman & Blake, 2002; Jastorff & Orban, 2009; Peelen, Wiggett & Downing, 2006; Peuskens, et al., 2005; Grezes et al., 2001; Vangeneugden et al., 2014), in addition to superior temporal sulcus (STS) as the integrative hub of encoding these visual features (Felleman & Van Essen, 1991; Jastorff et al., 2012; Oram & Perrett, 1996; Shiffrar, 1994; Vangeneugden et al., 2011; Vangeneugden, Pollick & Vogels, 2008) to eventually create action representations (Pitcher & Ungerleider, 2021); and finally parietal and frontal regions such as inferior parietal lobule (IPL), anterior intraparietal sulcus (aIPS), inferior frontal gyrus (IFG) and premotor cortex (PMC) that are important for processing higher aspects of observed actions leading to action understanding (Saygin, 2013; Rizzolatti et al., 1996; Rizzolatti & Craighero, 2004; Urgen & Orban, 2021; de C. Hamilton & Grafton, 2007; Jastorff et al., 2010; Buccino et al., 2001; Wheaton et al., 2004).

This network of regions is consistently evident in the literature that studied the BM stimuli under selective attention (Saygin et al., 2004; Herrington et al., 2012; Beauchamp, et al., 2003; Vaina, et al., 2001; Thornton, 2013; Grossman & Blake, 2002; Jastorff & Orban, 2009; Peelen et al., 2006). By definition, selective attention is our ability to focus on the task at hand while filtering out the task-irrelevant stimuli. Accordingly, BM perception under selective attention is studied with *active tasks* on BM stimuli such as discrimination based on walking direction, detection of BM under changing contrast, noise mask, or when the motion or form features are disrupted (e.g. inverted stimulus) or have become ambiguous (e.g. opposite motion of local versus global motion of BM stimulus) (Thornton, 2013). In such studies, the focus of attention is right at the BM stimulus itself such that top-down resources are available for BM perception (e.g. in the case of ambiguous stimuli). However, considering the innate tendency and social interference ability of humans towards the BM stimuli, BM perception can occur automatically in an effortless manner (Johansson, 1973; Thompson & Parasuraman, 2012; Thornton, Rensink & Shiffrar, 2002) even when the attention is directed away from it. Indeed, a behavioral experiment conducted by Thornton and Vuong (2004) showed that when presented as a task-irrelevant distractor at the periphery, BM impaired the performance on a task at the fovea. This finding shows that the “to-be-ignored” peripheral BM stimuli were processed incidentally. In such bottom-up processing, distinct regions in the BM perception network are expressed. In the case of passive viewing tasks, the activity maps include the visual feature encoding areas (i.e. EBA, hMT+, FBA) up until STS rather than the aforementioned BM processing areas that reach until fronto-parietal regions. For instance, in the study conducted by Jastorff and Orban (2009), only the occipital and ventral temporal areas (i.e. EBA, hMT+) showed significant activation when participants did not engage in an active task but just passively viewed the BM stimuli. Furthermore, Herrington and colleagues (2012) compared passive color detection versus active goal understanding task on BM stimuli. They found that right anterior STS, as well as hMT+ and EBA activations were stronger in goal understanding compared to the color detection task. Finally, in a retinotopic mapping study, Saygin & Sereno, 2008 showed that the spatial extent of retinotopic maps differed depending on whether the participants were engaged in an active task (attention to BM stimuli) or passive task (ignore BM stimuli). When attention is directed towards the BM stimuli on the wedge that swaps the visual field, retinotopic maps were observed in OTC including STS; parietal cortex including IPS and PPC; and frontal cortex including superior precentral sulcus (i.e. frontal eye field [FEF]). However, during the passive viewing task when attention is directed away from the BM stimuli, the retinotopic maps found in all these regions were either reduced or diminished to only OTC regions (V3, hMT+, LOC, STS) and IPS.

Considering the incidental processing of BM and the difference between its underlying neural correlates of top-down and bottom-up perception, one may hypothesize whether more “high-level” regions are affected by selective attention; whereas, feature encoding regions are involved in a rather “automatic” processing. Following this research question, in this study, we examined whether a stimulus that is socially and ecologically important like BM would be processed within the BM perception network when it was shown away from the focus of attention via an attentional load paradigm (Lavie, 1995).

In attentional load paradigms, an irrelevant stimulus is presented at the periphery, while participants are engaged in a visual detection task that demands either high or low attention at the fovea. Load theory proposed by Lavie (1995, 2005) suggests that considering there is a limit to perceptual capacity, the processing of each element within the field of view cannot be processed in the same manner. Attention, in that sense, works as a regulatory mechanism based on task demands (Bruckmaier et al., 2020). Accordingly, the activation related to the unattended, task-irrelevant stimuli is stronger under low attentional load since there are available attentional resources that can be allocated to these stimuli, while it is decreased or diminished under high load since attentional resources are limited. In the literature, flickering checkerboards, optic flow of lines, and colorful patches have been used as task-irrelevant stimuli on the periphery. The processing of these stimuli were found to be modulated by attentional load at early visual cortex areas (V1-V4), motion sensitive middle temporal (MT) cortex, and color responsive area V4, respectively (see Schwartz et al., 2005; Rees, 1997; Desseilles et al., 2009). By examining whether BM will be processed at the periphery through an attentional load experiment, this study is first to show the attentional load modulation on a socially meaningful, high-level stimulus.

By applying the attentional load paradigm, our aim is to answer two questions: (1) whether BM would be processed when it is shown away from the focus of attention as a task-irrelevant peripheral stimulus, and (2) if so, whether BM processing would be modulated by attentional load. By answering these questions, we can show (1) if BM is automatically processed in a bottom-up manner in the BM network, even when selective attention is not directed towards it; and (2) whether this bottom-up processing is modulated by a top-down factor that is attentional load. Thus, with this study design, we can examine the interaction between bottom-up and top-down processing of BM in a single task.

Considering the previous literature on the neural correlates of BM processing (Jastorff & Orban, 2009; Saygin & Sereno, 2008; Thompson & Parasuraman, 2012), we have hypothesized that:

1. BM stimulus would be processed in a bottom-up manner even if it was shown away from the focus of attention.
2. Visual feature encoding regions (i.e. hMT+, EBA) and possibly the higher level regions such as STS and fronto-parietal cortex areas (i.e. IPS, PMC) would be activated while processing BM incidentally.
3. The activation related to BM stimuli would be modulated by attentional load.

## 2. Methods

### 2.1. Participants

31 healthy volunteers (23 female; age range 18-31, mean age 23.05) with normal or corrected-to-normal visual acuity and no long-term use of medication for a neurological or psychiatric disorder participated in the study. The study was approved by the Human Research Ethics Committee of Bilkent University. Prior to the experiment, written informed consent and MRI prescreening form of each participant was collected. After the experiment, participants were compensated for their time.

### 2.2. Stimuli, Experimental Design and Procedure

There were two kinds of stimulus display presented onto the screen: (1) T-shaped stimuli shown at the fovea; (2) Point light displays (PLDs) of intact or scrambled BM stimulus shown at the periphery. All stimuli were generated and presented via the Psychophysics Toolbox (Psychtoolbox Version 3 [PTB-3]; Brainard, 1997; Pelli, 1997; Kleiner, et al., 2007) and BioMotion Toolbox (van Boxtel & Lu, 2013) on MATLAB (The Mathworks, Natick, MA).

The main task consisted of two pseudorandom rapid serial visual presentations (RSVP) of six t-shaped stimuli varying in their color (red, green or yellow) and orientation (upright or upside-down). The task was adapted from previous neuroimaging studies that examined attentional load (Bruckmaier et al., 2020; Desseilles et al., 2009; Rauss et al., 2012; Schwartz et al., 2005). A total of 12 t-shapes with four of them being the target stimuli were shown per condition for 500 ms each, comprising a total of 10 seconds with subsequent interstimulus intervals ranging between 250-350 ms (Figure 1.A). The number of target stimuli (four out of twelve) and overall t-shape displays were kept the same across blocks. Only the instruction screen differed based on the attentional load condition (Figure 1.B). Attentional load conditions differed as follows: in the low attentional load instruction, participants were asked to detect red shapes; meanwhile in the high attentional load, they had to detect upright yellow and upside-down green t-shapes. This manipulation is based on the study of Schwartz and colleagues (2005). Accordingly, detecting red shapes depends on the “pop-out” feature of one color among three others; meanwhile, detecting upright yellow or upside-down green t-shapes requires discrimination of a conjunction of color and orientation among all six combinations. Thus, the latter one demands more focused attention compared to the first one.

**Figure 1.**
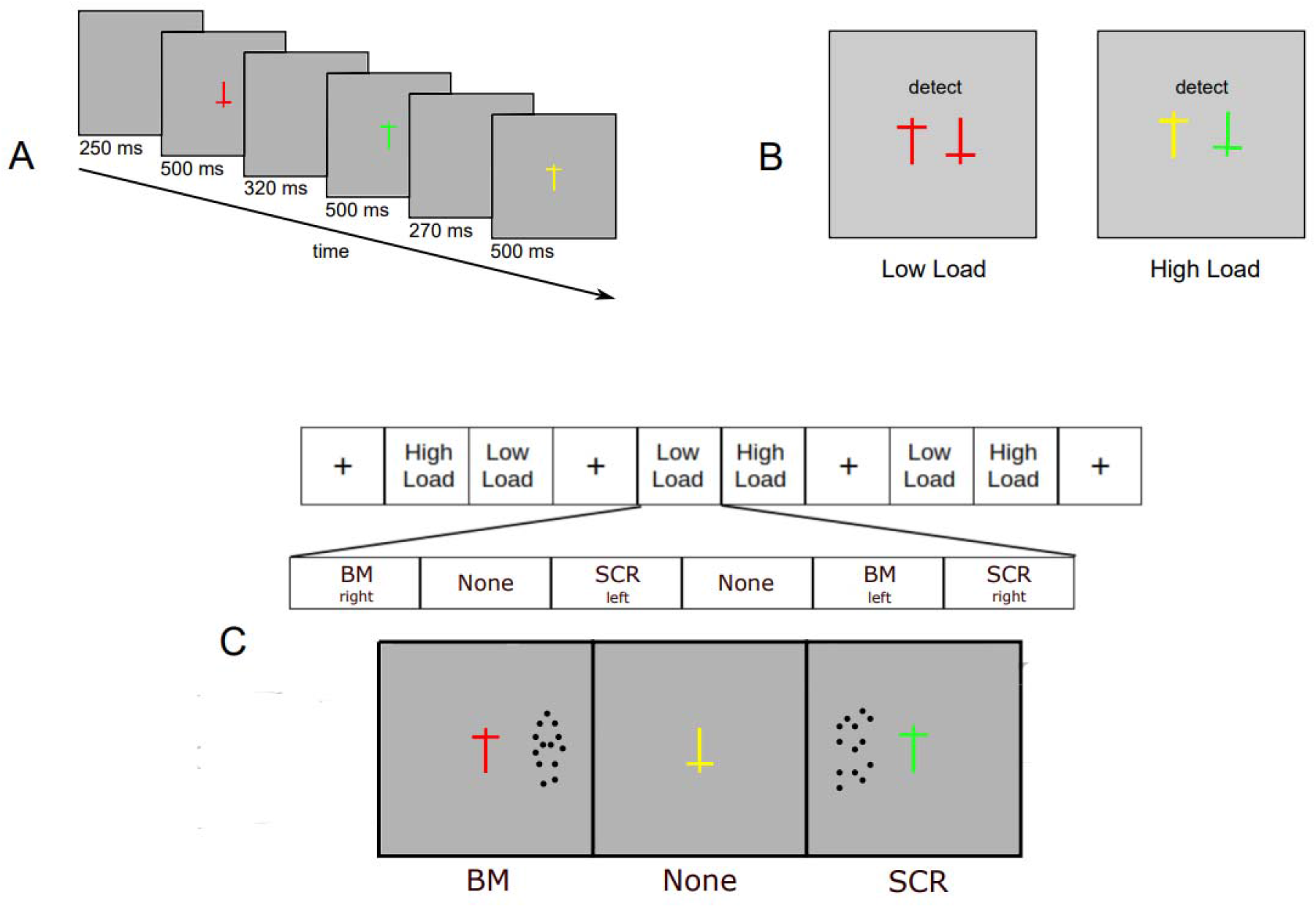
Stimuli design and procedure. Participants were engaged in a visual detection task at the fovea consisting of (A) t-shape like figures varying in their color (i.e. red, green, yellow) and orientation (i.e. upright, upside-down) shown for 500 ms with inter-stimulus intervals ranging between 250 and 350 ms randomly. (B) The main stimulus display was the same except for the instruction screen showing the target stimuli. (C) In each run, there were four rest and three pairs of experimental blocks. The display of the main task and the task-irrelevant peripheral stimulus (intact [BM] biological motion, scrambled [SCR] biological motion, or no stimulus at all [None] was shown in the periphery) was kept the same over each experimental block in all runs.

While participants were engaged in this visual detection task varying in attentional load, either intact or scrambled point-light displays of BM or no stimulus at all were shown 4 visual degrees away from the foveal t-shape stimuli (Figure 1.C). The PLD animation of a walking human was downloaded from an open motion capture database (CMU Graphics Lab, http://mocap.cs.cmu.edu/) and then edited on Motion Kinematic and Kinetic Analyser software (MOKKA, 2018, https://biomechanical-toolkit.github.io/mokka/) to acquire a 2D front view with a total of 13 dots comprising a head, shoulders, elbows, hands, waist bones, knees, and feet joint locations. Each animation was reduced to one second and consisted of 61 frames. In order to have the same peripheral stimuli throughout the RSVP of twelve t-shapes, one animation was shown on a total of 10 seconds-long loop. The scrambled point-light motion stimulus (SCR) was created via BioMotion Toolbox (van Boxtel & Lu, 2013) by shuffling the anchor positions of dots in the intact BM stimulus while keeping each dot’s local motion trajectories the same. Therefore, SCR had the same number of dots, and the same local dot motion as in intact BM but the global form and motion of the intact BM was disrupted in SCR.

The fMRI experiment consisted of 8 runs. There were four rest and three pairs of experimental blocks in each run. Each run started, ended and was interleaved with rest blocks in which the participant had to fixate on a cross shown at the fovea for 8, 10, 12, or 14 seconds (11 seconds on average). The experimental blocks were shown as subsequent pairs (Figure 1.C) with an instruction screen lasting 5 seconds prior to their start showing the target stimuli based on the attentional load condition. The order of blocks was counterbalanced. Each experimental block consisted of six peripheral stimulus conditions that were randomized in their order: BM-right, BM-left, SCR-right, SCR-left, No-peripheral-stimulus condition (None) (Figure 1.C). The None condition was shown twice to equal the number of conditions within a block.

Prior to scanning, participants were familiarized with the intact and scrambled BM stimuli separately via an explicit overview of each stimulus, and were tested whether they could discriminate them on the periphery during a fixation on a foveal cross as a separate practice. Lastly, the main task was practiced without any peripheral stimuli for 36 trials per load condition. After assuring a 80% success rate, participants were taken to the scanner room. During the fMRI experiment, they had to detect the target stimuli shown in the instructions as accurately as possible, and they were explicitly told to fixate on the t-shape stimulus and avoid looking at any other stimuli that may appear in the periphery. The stimuli were shown on an MR compatible LED screen (TELEMED, 60 Hz refresh rate, 800x600 pixel, 32 inch) and seen via a mirror mounted on the head-coil. Participants’ responses were collected via an MR suitable fiber optic button box.

### 2.3. Behavioral Data Analysis

A 2 x 3 within-subjects analysis of variance (ANOVA) was conducted (N=31) to test the effect of attentional load (AL: High, Low) and peripheral stimulus (PS: BM, SCR, None) on reaction times and accuracy. Threshold of p-value was set at p < 0.05 under Bonferroni correction.

### 2.4. MRI Data Acquisition and Analyses

Scanning took place at the National Magnetic Resonance Research Center (UMRAM) in Bilkent University with 3T Siemens TimTrio MR scanner with a 32-channel phase array head coil. In order to minimize head movement, participants’ heads were supported with foam padding. Before the experimental scans, high resolution T1-weighted structural images were obtained (TR=2600 ms, TE=2.92 ms, flip angle=12°, FoV read=256 mm, FoV phase=87.5%, 176 slices with 1×1×1 mm3 resolution). Then, over eight runs, 227 functional images were acquired using gradient-echo planar imaging (TR=2000 ms, TE=22 ms, flip angle=90°, 64 x 64 matrix, FoV read=192 mm, 43 slices with a thickness of 2.5mm, 3×3×2.5 mm3 resolution).

#### 2.4.1. Anatomical and Functional Data Preprocessing via fMRIPrep

Results included in this study come from preprocessing performed using fMRIPrep 20.1.1 (Esteban, Markiewicz, et al. (2018); Esteban, Blair, et al. (2018); RRID:SCR_016216), which is based on Nipype 1.5.0 (Gorgolewski et al. (2011); Gorgolewski et al. (2018); RRID:SCR_002502).

##### Anatomical data preprocessing

A total of 2 T1-weighted (T1w) images were found within the input BIDS dataset. All of them were corrected for intensity non-uniformity (INU) with N4BiasFieldCorrection (Tustison et al. 2010), distributed with ANTs 2.2.0 (Avants et al. 2008, RRID:SCR_004757). The T1w-reference was then skull-stripped with a Nipype implementation of the antsBrainExtraction.sh workflow (from ANTs), using OASIS30ANTs as target template. Brain tissue segmentation of cerebrospinal fluid (CSF), white-matter (WM) and gray-matter (GM) was performed on the brain-extracted T1w using fast (FSL 5.0.9, RRID:SCR_002823, Zhang, Brady, and Smith 2001). A T1w-reference map was computed after registration of 2 T1w images (after INU-correction) using mri_robust_template (FreeSurfer 6.0.1, Reuter, Rosas, and Fischl 2010). Brain surfaces were reconstructed using recon-all (FreeSurfer 6.0.1, RRID:SCR_001847, Dale, Fischl, and Sereno 1999), and the brain mask estimated previously was refined with a custom variation of the method to reconcile ANTs-derived and FreeSurfer-derived segmentations of the cortical gray-matter of Mindboggle (RRID:SCR_002438, Klein et al. 2017). Volume-based spatial normalization to one standard space (MNI152NLin2009cAsym) was performed through nonlinear registration with antsRegistration (ANTs 2.2.0), using brain-extracted versions of both T1w reference and the T1w template. The following template was selected for spatial normalization: ICBM 152 Nonlinear Asymmetrical template version 2009c [Fonov et al. (2009), RRID:SCR_008796; TemplateFlow ID: MNI152NLin2009cAsym],

##### Functional data preprocessing

For each of the 8 BOLD runs found per subject (across all tasks and sessions), the following preprocessing was performed. First, a reference volume and its skull-stripped version were generated using a custom methodology of fMRIPrep. Head-motion parameters with respect to the BOLD reference (transformation matrices, and six corresponding rotation and translation parameters) are estimated before any spatiotemporal filtering using mcflirt (FSL 5.0.9, Jenkinson et al. 2002). BOLD runs were slice-time corrected using 3dTshift from AFNI 20160207 (Cox and Hyde 1997, RRID:SCR_005927). Susceptibility distortion correction (SDC) was omitted. The BOLD reference was then co-registered to the T1w reference using bbregister (FreeSurfer) which implements boundary-based registration (Greve and Fischl 2009). Co-registration was configured with six degrees of freedom. The BOLD time-series (including slice-timing correction when applied) were resampled onto their original, native space by applying the transforms to correct for head-motion. These resampled BOLD time-series will be referred to as preprocessed BOLD in original space, or just preprocessed BOLD. The BOLD time-series were resampled into standard space, generating a preprocessed BOLD run in MNI152NLin2009cAsym space. First, a reference volume and its skull-stripped version were generated using a custom methodology of fMRIPrep. Several confounding time-series were calculated based on the preprocessed BOLD: framewise displacement (FD), DVARS and three region-wise global signals. FD was computed using two formulations following Power (absolute sum of relative motions, Power et al. (2014)) and Jenkinson (relative root mean square displacement between affines, Jenkinson et al. (2002)). FD and DVARS are calculated for each functional run, both using their implementations in Nipype (following the definitions by Power et al. 2014). The three global signals are extracted within the CSF, the WM, and the whole-brain masks. Additionally, a set of physiological regressors were extracted to allow for component-based noise correction (CompCor, Behzadi et al. 2007). Principal components are estimated after high-pass filtering the preprocessed BOLD time-series (using a discrete cosine filter with 128s cut-off) for the two CompCor variants: temporal (tCompCor) and anatomical (aCompCor). tCompCor components are then calculated from the top 5% variable voxels within a mask covering the subcortical regions. This subcortical mask is obtained by heavily eroding the brain mask, which ensures it does not include cortical GM regions. For aCompCor, components are calculated within the intersection of the aforementioned mask and the union of CSF and WM masks calculated in T1w space, after their projection to the native space of each functional run (using the inverse BOLD-to-T1w transformation). Components are also calculated separately within the WM and CSF masks. For each CompCor decomposition, the k components with the largest singular values are retained, such that the retained components’ time series are sufficient to explain 50 percent of variance across the nuisance mask (CSF, WM, combined, or temporal). The remaining components are dropped from consideration. The head-motion estimates calculated in the correction step were also placed within the corresponding confounds file. The confound time series derived from head motion estimates and global signals were expanded with the inclusion of temporal derivatives and quadratic terms for each (Satterthwaite et al. 2013). Frames that exceeded a threshold of 0.5 mm FD or 1.5 standardized DVARS were annotated as motion outliers. All resamplings can be performed with a single interpolation step by composing all the pertinent transformations (i.e. head-motion transform matrices, susceptibility distortion correction when available, and co-registrations to anatomical and output spaces). Gridded (volumetric) resamplings were performed using antsApplyTransforms (ANTs), configured with Lanczos interpolation to minimize the smoothing effects of other kernels (Lanczos 1964). Non-gridded (surface) resamplings were performed using mri_vol2surf (FreeSurfer).

Many internal operations of fMRIPrep use Nilearn 0.6.2 (Abraham et al. 2014, RRID:SCR_001362), mostly within the functional processing workflow. For more details of the pipeline, see <u>the section corresponding to workflows in fMRIPrep’s documentation</u>.

### 2.5. Univariate Analysis and Activation Maps

After preprocessing, first- and second-level analyses were conducted via SPM12 software package (http://www.fil.ion.ucl.ac.uk/spm/software/spm12/) on MATLAB (The Mathworks, Natick, MA). A design matrix of fourteen regressors (six experimental conditions: BM under High Load [HBM], BM under Low Load [LBM], SCR under High Load [HSCR], SCR under Low Load [LSCR], None under High Load [HNone], None under Low Load [LNone]; one fixation block; six motion regressors: three translations and three rotations; one constant factor) was created for the first-level general linear model (GLM). Each regressor was convolved with the default canonical hemodynamic response function on SPM12. Following the first-level analysis, group level analysis was performed as a 2 (AL: High, Low) x 3 (PS: BM, SCR, None) within-subjects flexible factorial design (Penny et al., 2011). In order to run flexible factorial analysis, subjects and experimental conditions (AL and PS) were defined as independent and dependent factors respectively. No additional parameter or covariant was added. Further linear contrasts tested in the group analysis were based on understanding the main effect of AL and PS conditions separately. Activation maps were thresholded at p < 0.05 under familywise error (FWE) correction.

### 2.6. Searchlight-based Multivariate Pattern Analysis (Decoding)

In order to discriminate the pattern of brain activity associated with different attentional load modulations (High versus Low) and peripheral stimuli conditions (BM, SCR, None), several decoding analyses were done. Each of six experimental conditions was labeled accordingly in each classification: HBM, HSCR, HNone, LBM, LSCR, LNone. In order to differentiate the pattern of activation in high and low AL blocks, “High versus Low” binary classification (Supplementary Figure 1) was run. Furthermore, three-way (Supplementary Figure 2) and binary PS classifications were run on PS classes to observe the difference of activation patterns associated with each PS condition: BM versus SCR versus None; BM versus None; SCR versus None; BM versus SCR. Further binary classifications were run in order to discriminate the attentional load modulation on each peripheral stimulus condition to be compared with each other: HBM versus HNone, LBM versus LNone; HSCR versus HNone, LSCR versus LNone; HBM versus HSCR, LBM versus LSCR. Whole-brain decoding maps were created through a searchlight-based (3 mm radius sphere) multivariate pattern analysis (MVPA) with a linear support vector machine (LibSVM; regularization parameter C = 1) on the Decoding Toolbox (TDT; Hebart, Görgen & Haynes, 2015). Runwise beta images were extracted from the first-level GLM of each subject and used as inputs for the classifiers. Leave-one-run-out cross-validation procedure was applied and the accuracy minus chance brain images of each subject were created to be used in the one-sample t-test group analysis. The applied threshold was at p < 0.05 under FWE correction.

## 3. Results

### 3.1 Behavioral Results

#### Accuracy

The task was completed successfully by participants as indicated by an overall accuracy being more than 80 percent. There was a main effect of attentional load (F(1,30) = 85.307, p <0.001, η^2^ = 0.671). The accuracy rate in the low load (M = 0.97, SD = 0.03) was significantly higher than that of high load (M = 0.84, SD = 0.081) (t(30)= 9.236, p <0.001). However, there was no main effect of PS ((F(1,30) = 1.231, p=0.299), and no interaction between the load and the PS conditions ((F(1,30) = 2.071, p=0.135). The results showed that regardless of the PS, the AL effect was evident on the accuracy rates.

#### Reaction Times

Reaction time (RT) analysis showed that there was a main effect of both AL (F(1,30) = 82.507,p <0.001, η^2^ = 0.33) and PS (F(1,30) = 115.956, p <0.001, η^2^ = 0.273) (Table 1). The RTs found in the high load was significantly higher than that of low load (t(30)= 8.897, p <0.001). Post-hoc tests on the PS conditions showed that participants’ RTs during no PS condition was significantly higher than that of BM (t(30) = 12.976, p <0.001) and SCR (t(30) = 13.391, p <0.001) conditions. While, the RTs in intact and scrambled BM conditions were not significantly different from each other (t(30) = -1.196, p=0.709).

**Table 1.**
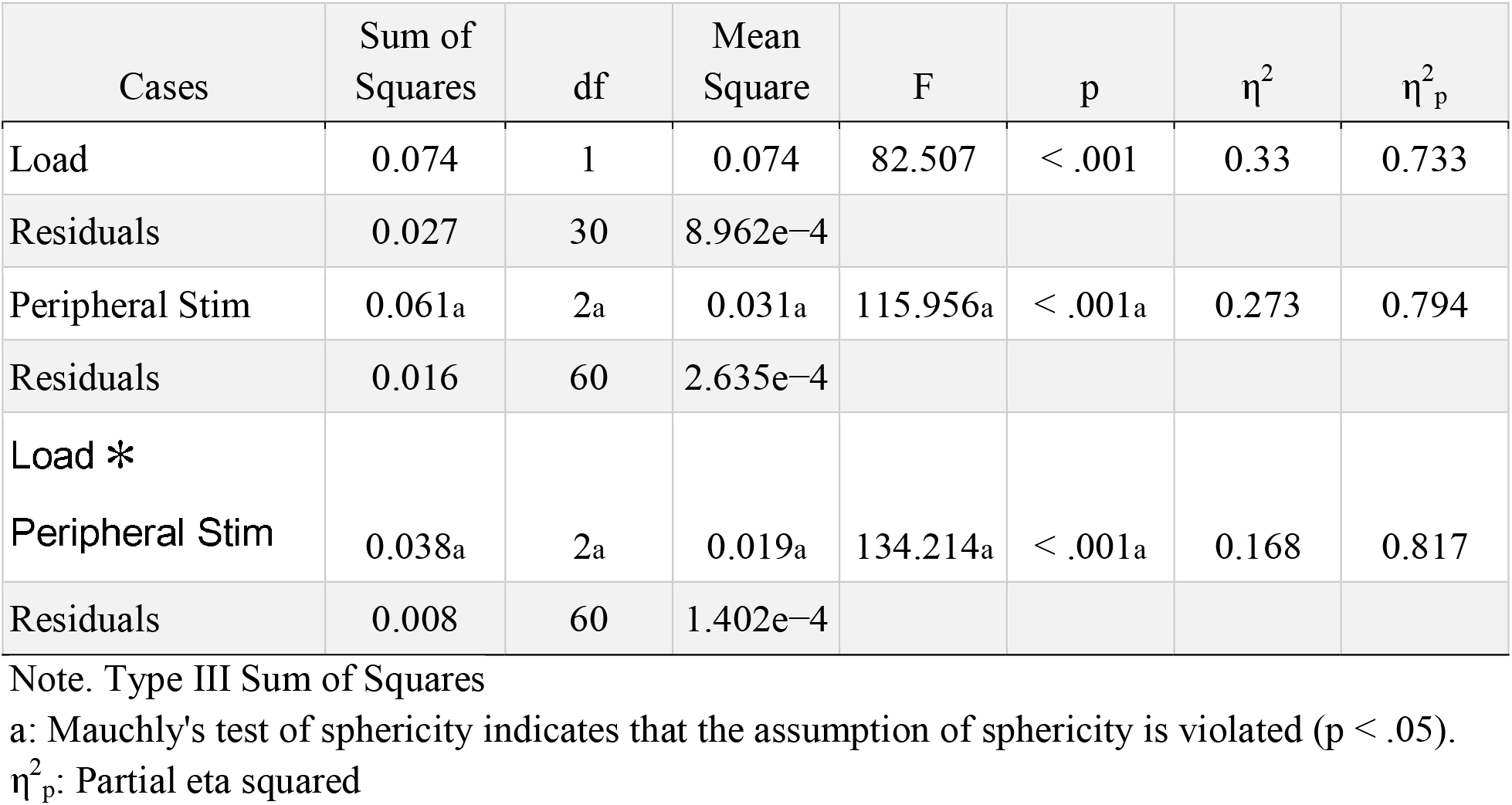
Within Subjects Effects

Moreover, a significant interaction between the load and PS was also found (F(1,30) = 134.214, p <0.001, η2 = 0.168). Post-hoc tests on each of six conditions revealed that although there was a significant difference between SCR versus None (HSCR > HNone: t(30) = - 19.490, p <0.001) and BM versus None conditions (HBM > HNone: t(30) = - 18.497, p <0.001), SCR and BM conditions did not significantly differ from each other under the high load (HBM > HSCR: t(30)= 0.993, p = 1). None of the pair-wise PS comparisons were significantly different from each other under low attentional load comparisons (Table 2).

**Table 2.**
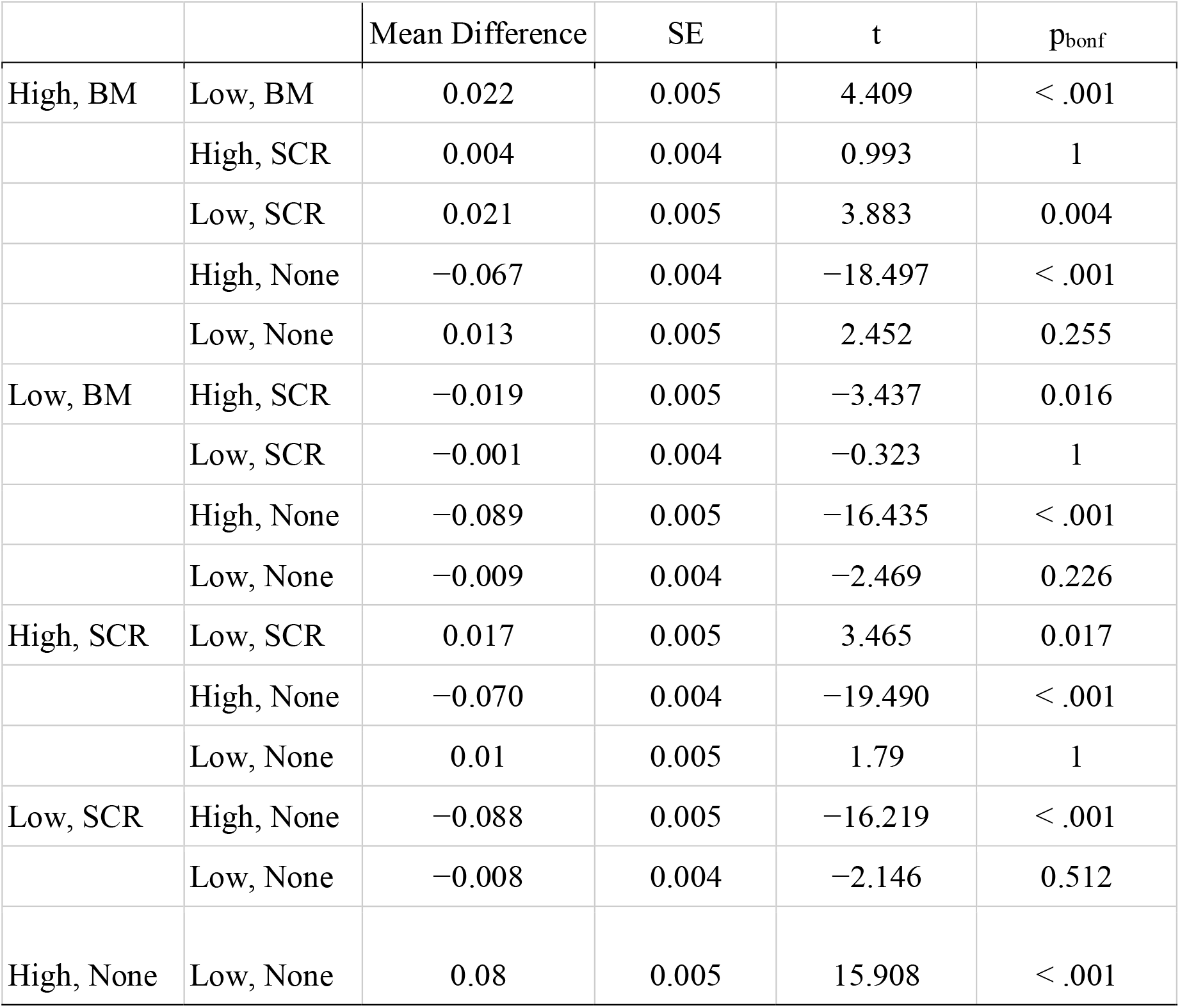
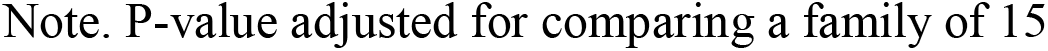
Post Hoc Comparisons - Load ✻ Peripheral Stim

|  |  | Mean Difference | SE | t | p<sub>bonf</sub> |
| --- | --- | --- | --- | --- | --- |
| High, BM | Low, BM | 0.022 | 0.005 | 4.409 | < .001 |
|  | High, SCR | 0.004 | 0.004 | 0.993 | 1 |
|  | Low, SCR | 0.021 | 0.005 | 3.883 | 0.004 |
|  | High, None | -0.067 | 0.004 | -18.497 | < .001 |
|  | Low, None | 0.013 | 0.005 | 2.452 | 0.255 |
| Low, BM | High, SCR | -0.019 | 0.005 | -3.437 | 0.016 |
|  | Low, SCR | -0.001 | 0.004 | -0.323 | 1 |
|  | High, None | -0.089 | 0.005 | -16.435 | < .001 |
|  | Low, None | -0.009 | 0.004 | -2.469 | 0.226 |
| High, SCR | Low, SCR | 0.017 | 0.005 | 3.465 | 0.017 |
|  | High, None | -0.070 | 0.004 | -19.490 | < .001 |
|  | Low, None | 0.01 | 0.005 | 1.79 | 1 |
| Low, SCR | High, None | -0.088 | 0.005 | -16.219 | < .001 |
|  | Low, None | -0.008 | 0.004 | -2.146 | 0.512 |
| High, None | Low, None | 0.08 | 0.005 | 15.908 | < .001 |

Note. P-value adjusted for comparing a family of 15

Overall, behavioral results revealed that the attentional load effect was evident on each PS condition. Participants responded faster when there was a task-irrelevant stimulus (BM or SCR) in the periphery as compared to nothing during high attentional load; but there was no difference between the intact and scrambled BM conditions in neither high nor low attentional load blocks.

### 3.2. Whole-brain Univariate Activation Maps

In order to examine the whole-brain activity maps during the bottom-up perception of BM, we have conducted whole-brain within-subject flexible factorial (FF) analysis. All results are thresholded at FWE-corrected, p < 0.05; and the peak-level results are reported.

#### Main effect of Attentional Load

The main effect of AL as indicated by an F-test on high and low AL blocks yielded two distinct network activity: Fronto-parietal attention network and default mode network (DMN) (Table 3, Figure 2.a).

**Figure 2.**
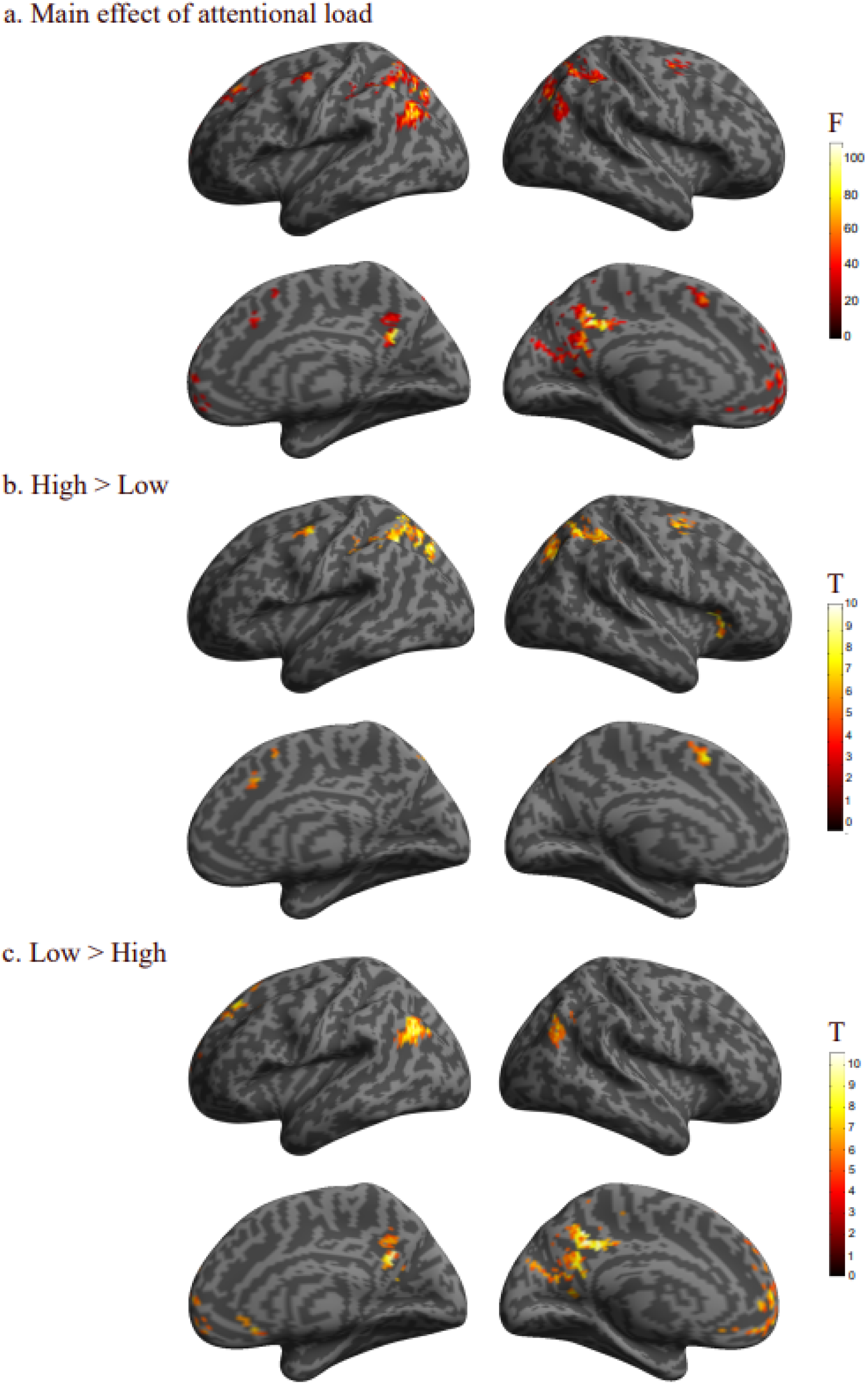
Univariate activation maps for attentional load. (a) The activity maps of the main effect of attentional load. The F-test results showed fronto-parietal attention and DMN region activations. (b) The activity maps of the High >Low attentional load t-contrast. The t-contrast of High >Low showed fronto-parietal network activation. (c) The activity maps of the Low >High attentional load t-contrast. The t-contrast of Low >High showed activation within DMN. Results were FWE-corrected, at a threshold of p <.05.

**Table 3.** Main effect of attentional load. Results were FWE-corrected, at a threshold of p <.05.

| **Main Effect of Attentional Load** |  |  |
| --- | --- | --- |
| *Region* | *Coordinates* | *F-Score* |
| L Precuneus | -4 -55 19 | 127.55 |
| L Angular Gyrus (IPL) (BA23) | -46 -79 37 | 115.7 |
| L Middle Orbital Gyrus (BA39) | -1 60 -11 | 108.52 |
| L Inferior Parietal Lobule (BA39) | -25 -70 42 | 104.65 |
| L Superior Frontal Gyrus (BA6) | -31 -4 69 | 86.81 |
| L Posterior-Medial Frontal (BA6) | -4 9 52 | 81.87 |
| R Middle Occipital Gyrus (BA7) | 30 -67 32 | 85.15 |
| R Angular Gyrus (IPL) (BA39) | 51 -73 34 | 74.16 |
| R Middle Frontal Gyrus (BA6) | 27 -7 49 | 69.01 |
| **High > Low** |  |  |
| *Region* | *Coordinates* | *T-Score* |
| L Inferior Parietal Lobule (BA39) | -25 -70 42 | 10.23 |
| L Superior Frontal Gyrus (BA6) | -31 -4 69 | 9.32 |
| L Posterior-Medial Frontal (BA6) | -4 9 52 | 9.05 |
| R Middle Occipital Gyrus (BA7) | 30 -67 32 | 9.23 |
| R Insula Lobe (BA45) | 33 24 9 | 8.65 |
| R Middle Frontal Gyrus (BA6) | 27 -7 49 | 8.31 |
| **Low > High** |  |  |
| *Region* | *Coordinates* | *T-Score* |
| L Precuneus (BA23) | -4 -55 19 | 11.29 |
| L Angular Gyrus (IPL) (BA39) | -46 -79 37 | 10.76 |
| L Middle Orbital Gyrus (BA10) | -1 60 -11 | 10.42 |
| R Angular Gyrus (IPL) (BA39) | 51 -73 34 | 8.61 |

Further t-contrast analysis revealed that for the High > Low contrast, in line with the literature (Chong et al., 2008: Desseilles et al., 2009: Schwartz et al., 2004), significant activation was found within the left inferior parietal lobule (peak: -25 -70 42; T= 10.23) and left superior frontal gyrus (peak: -31 -4 69; T= 9.32) that constitute the fronto-parietal attention network (Figure 2.b). Additionally, posterior medial frontal gyrus in the left hemisphere that is associated with error-monitoring and post-error adjustments (Danielmeier et al., 2011) were found activated (peak= -4 9 52; T= 9.05). For the Low > High t-contrast, DMN regions (Raichle, 2015) were observed (Figure 2.c), revealing activity maps in left precuneus (peak: -4 -55 19; T= 11.29), left and right angular gyri (peak: -46 -79 37; F=10.76) (peak: 51 -73 34; T= 8.61), and left middle orbital gyrus (peak= -1 60 -11; T= 10.42) (Table 3).

#### Main effect of Peripheral Stimulus

The main effect of PS as indicated by an F-test on BM, SCR, and None conditions yielded bilateral activity maps in secondary visual (BA18) and visual association (BA19) cortices including motion sensitive MT/V5 region as expected (Figure 3.a, Table 4).

**Figure 3.**
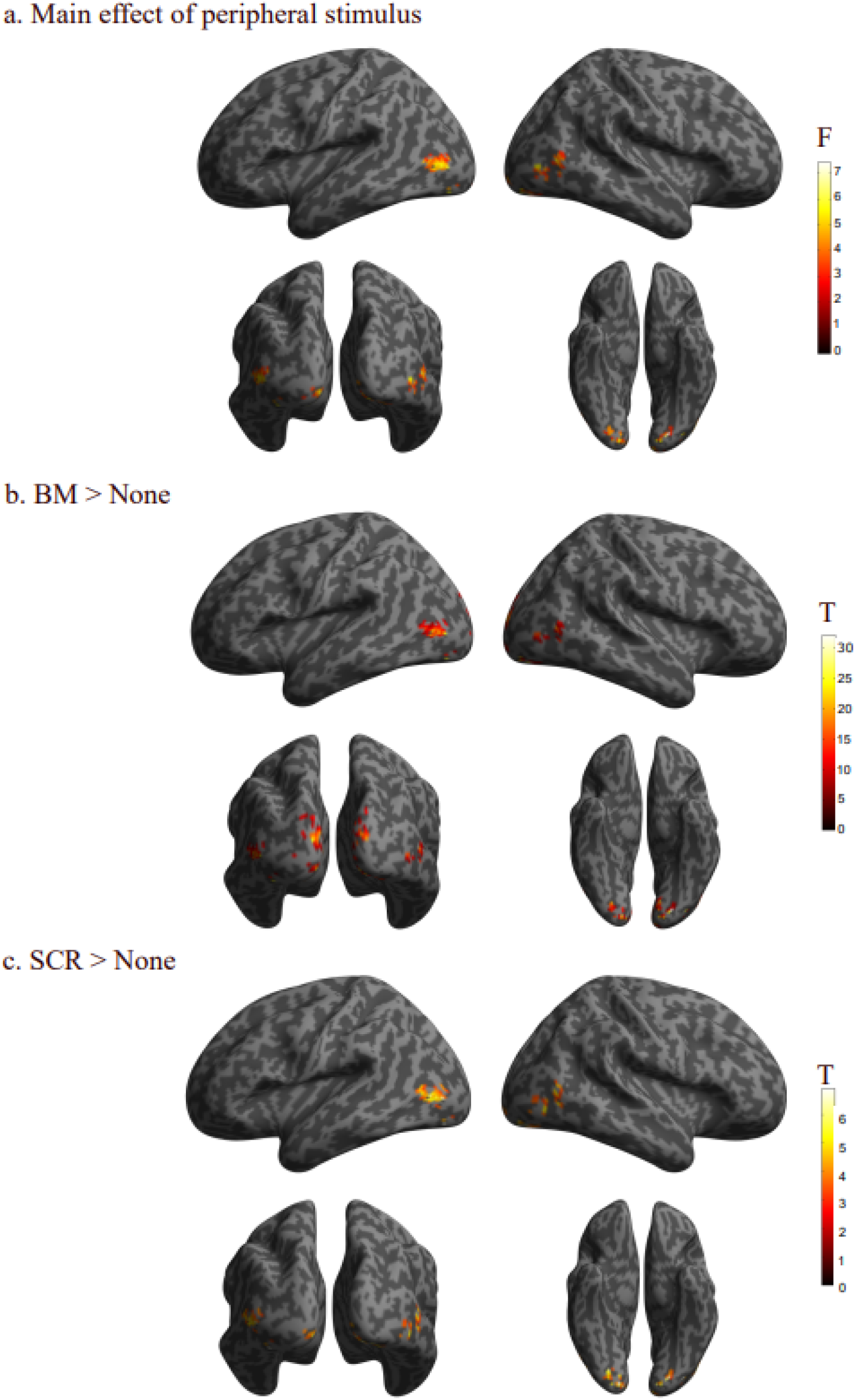
Univariate activation maps for peripheral stimulus. (a) The activity maps of the main effect of peripheral stimulus. The F-test results showed activity maps in medial early visual cortex and motion-sensitive regions in the OTC. (b) The activity maps of the BM >None t-contrast. The t-contrast of BM >None showed activation at motion-sensitive regions in the OTC. (c) The activity maps of the SCR >None t-contrast. The t-contrast of SCR >None yielded activation at motion-sensitive regions in the OTC. Results were at an uncorrected threshold of p <.001.

**Table 4.**
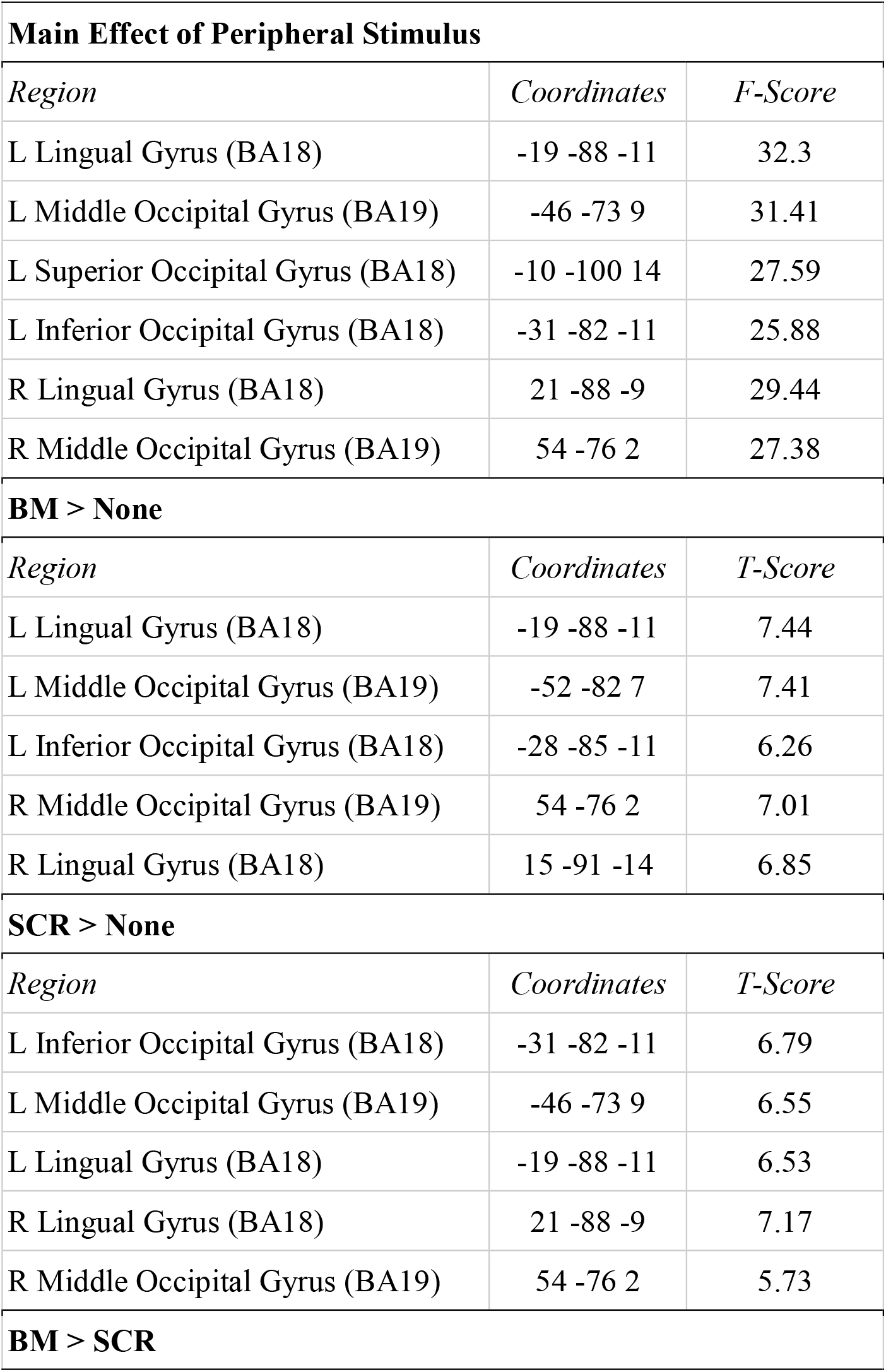
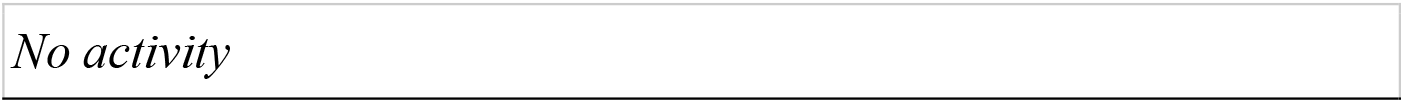
Main effect of peripheral stimulus. Results were at an uncorrected threshold of p <.001.

Furthermore, separate comparisons between each PS and None conditions yielded similar activities for both BM > None and SCR > None t-contrasts (Table 4): bilateral secondary visual (BA18) and visual association (BA19) cortices including motion-sensitive MT/V5 regions were activated in both contrasts (Figure 3.b).

Overall, the bottom-up processing of PS was observed in motion-sensitive visual areas regardless of the attentional load. Contrary to our expectation, there was no difference between intact and scrambled BM stimuli.

#### Attentional Load Modulation on Peripheral Stimulus

AL modulation on PS is indicated by the decreased activation or diminished activity maps in high load blocks as compared to low load blocks. Accordingly, while there was a significant activation found within the OTC regions towards the BM > None and SCR > None contrasts under low load, only an early visual cortex region showed activity towards the same contrasts under high load (Table 5).

**Table 5.**
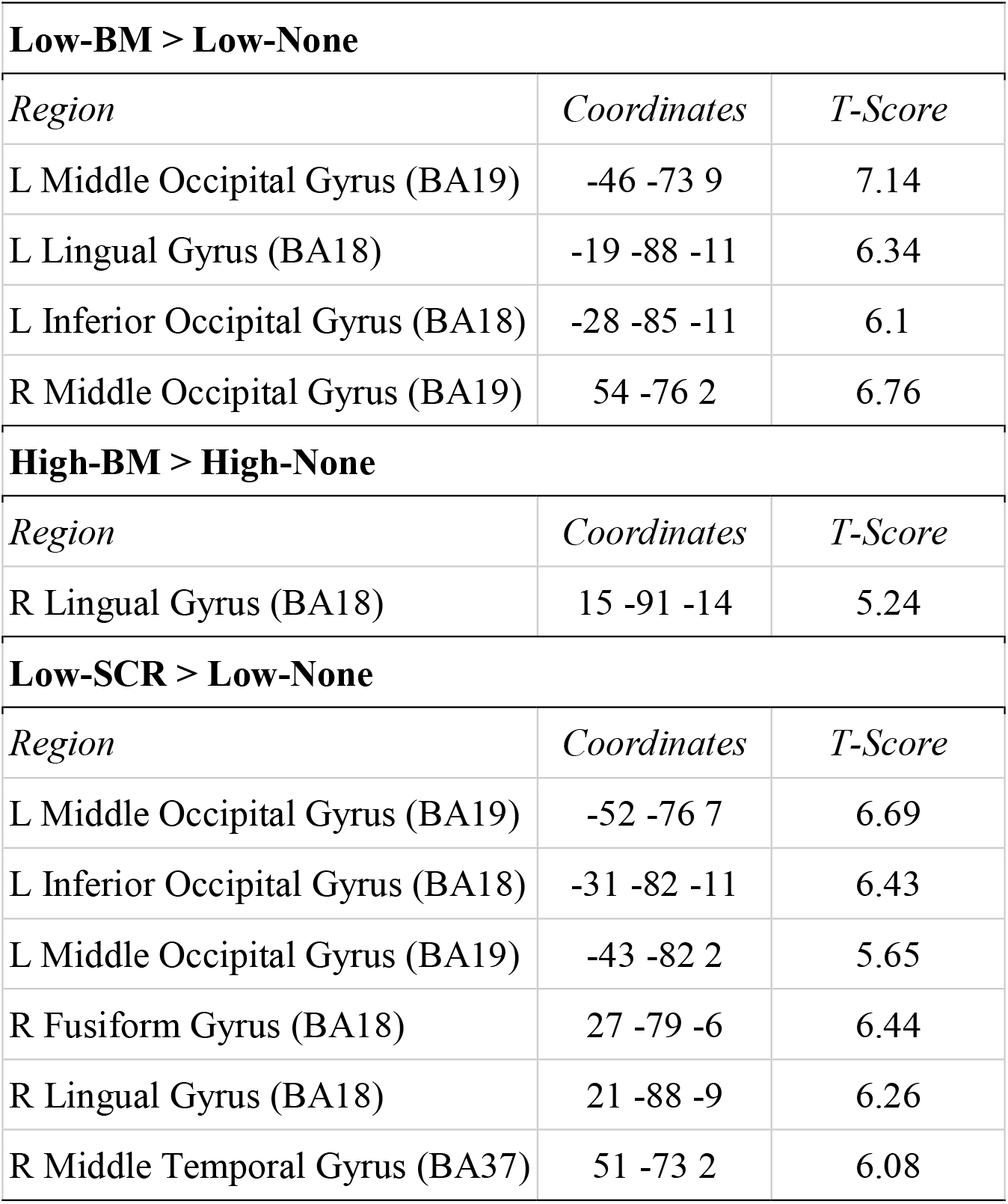
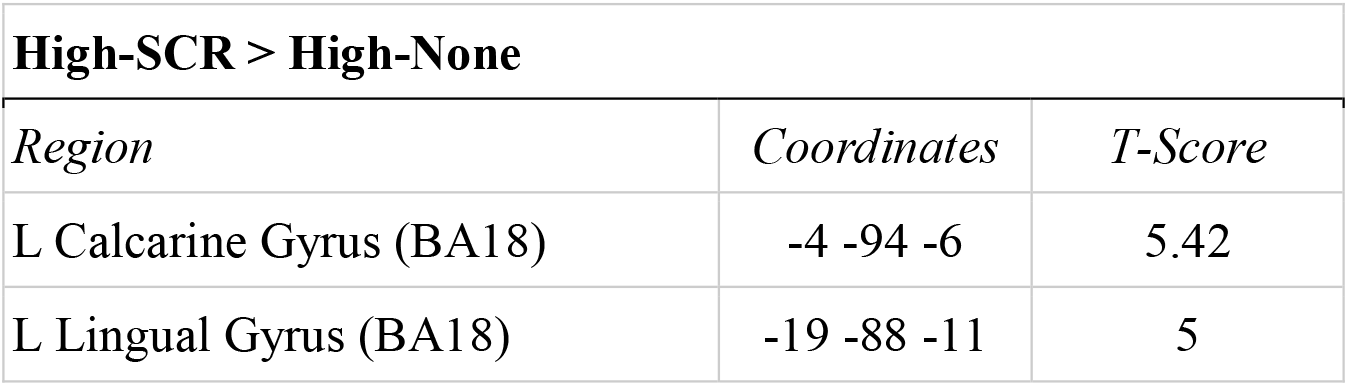
Attentional load modulation on peripheral stimulus. Results were at an uncorrected threshold of p <.001.

### 3.3. Whole-brain Searchlight based MVPA

Considering the increased sensitivity of multivariate pattern analysis compared to univariate analysis (Davis et al., 2014), we conducted a searchlight based MVPA on the whole-brain to test which brain regions could discriminate between the processing of each PS condition under high and low attentional load. The group-level one sample t-test results on chance minus accuracy brain images of each participant was calculated for the following binary classifications: BM vs None, LBM vs LNone, HBM vs HNone, SCR vs None, LSCR vs LNone, HSCR vs HNone.

#### BM vs None Classification Results

Regardless of the attentional load block, motion and form sensitive regions in OTC could significantly decode BM and None conditions (Figure 4.a). The AL modulation was evident as indicated by the diminished extent and amount of the regions that could discriminate between the BM and None classes under the high load as compared to low load (Figure 4.b and Figure 4.c). Specifically, the decoding regions for BM vs None were at peak in the bilateral middle occipital gyrus (Left: peak: -46 -82 4; T= 9.92; Right: peak: 42 -73 4; T= 7) and left lingual gyrus (peak: - 16 -88 -14; T= 15.4). For the same comparison under low attentional load, the regions that could significantly separate LBM and LNone classes included calcarine gyrus (peak: -13 -97 -4; T= 15.51) and middle occipital gyrus (peak: -49 -79 2; T= 10.5) in the left hemisphere, in addition to the lingual gyrus (peak: 24 -85 -14; T= 7.13) in the right hemisphere. As expected, due to the modulation of attentional load, the same comparison under high load showed only calcarine gyrus in the left hemisphere (peak: -10 -91 -4; T= 13.59) as the region that significantly separates HBM and HNone from each other (Table 6).

**Figure 4.**
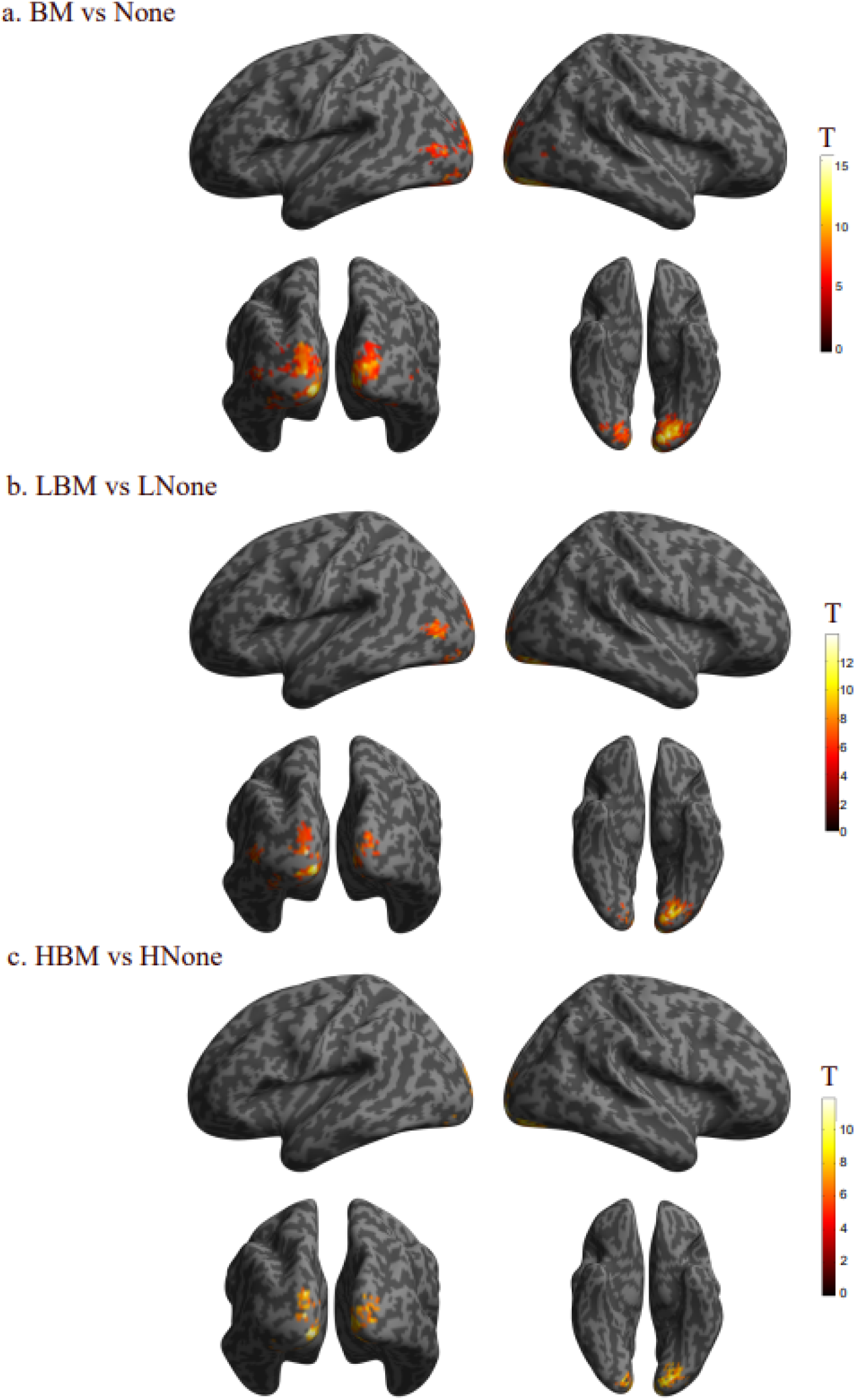
Multivariate pattern analysis results of the attentional load modulation on intact BM stimuli. (a) Decoding regions of the BM versus None classification: Motion and form sensitiv OTC regions were shown to discriminate BM and None conditions regardless of the attentional load. (b) Attentional load modulation on BM was evident: Under low attentional load, BM and None could be decoded in the OTC regions; while (c) under high load, only the early visual areas could discriminate BM and None stimuli. Results were FWE-corrected, at a threshold of p <.05.

**Table 6.** Attentional load modulation on BM indicated by the diminished extent of decoding regions. Results were FWE-corrected, at a threshold of p <.05.

| **BM vs None** |  |  |
| --- | --- | --- |
| *Region* | *Coordinates* | *T-Score* |
| L Lingual Gyrus (BA18) | -16 -88 -14 | 15.4 |
| L Middle Occipital Gyrus (BA19) | -46 -82 4 | 9.92 |
| R Middle Occipital Gyrus (BA19) | 42 -73 4 | 7 |
| **Low-BM vs Low-None** |  |  |
| *Region* | *Coordinates* | *T-Score* |
| L Calcarine Gyrus (BA18) | -13 -97 -4 | 15.51 |
| L Middle Occipital Gyrus (BA19) | -49 -79 2 | 10.5 |
| R Lingual Gyrus (BA19) | 24 -85 -14 | 7.13 |
| **High-BM vs High-None** |  |  |
| *Region* | *Coordinates* | *T-Score* |
| L Calcarine Gyrus (BA18) | -10 -91 -4 | 13.59 |
| **SCR vs None** |  |  |
| *Region* | *Coordinates* | *T-Score* |
| L Fusiform Gyrus (BA18) | -22 -82 -11 | 14.32 |
| L Middle Occipital Gyrus (BA19) | -49 -76 14 | 9.38 |
| L Middle Occipital Gyrus (BA19) | -31 -70 -6 | 6.79 |
| **Low-SCR vs Low-None** |  |  |
| *Region* | *Coordinates* | *T-Score* |
| L Calcarine Gyrus (BA18) | -10 -94 -4 | 12.13 |
| L Superior Occipital Gyrus (BA18) | -19 -97 17 | 9.06 |
| L Middle Occipital Gyrus (BA19) | -52 -82 4 | 8.67 |
| L Middle Occipital Gyrus (BA19) | -40 -88 4 | 7.12 |
| **High-SCR vs High-None** |  |  |
| *Region* | *Coordinates* | *T-Score* |
| L Calcarine Gyrus (BA18) | -7 -91 -6 | 14.85 |
| L Superior Occipital Gyrus (BA18) | -13 -97 19 | 9.28 |
| L Superior Occipital Gyrus (BA18) | -13 -97 29 | 6.91 |

#### SCR vs None Classification Results

The SVM could discriminate between the SCR and None conditions over all AL blocks at the motion and form sensitive OTC regions (Figure 5.a). The modulation of AL was evident as indicated by the decreased amount and extension of decoding regions in the high load as compared to that of low load (Figure 5.b and Figure 5.c). Specifically, regardless of the AL block, SCR versus None classification could be decoded at fusiform gyrus (peak: -22 -82 -11; T= 14.32) and middle occipital gyrus (peak: -49 -76 14; T= 9.38; peak: -31 -70 -6; T= 6.79) in the left hemisphere. Moreover, under the low load, SCR and None conditions could be discriminated significantly at the calcarine gyrus (peak: -10 -94 -4; T= 12.13), superior occipital gyrus (peak: - 19 -97 17; T= 9.06), and middle occipital gyrus (peak: -52 -82 4, T= 7.12) regions in the left hemisphere. Under high load, the same comparison of SCR and None conditions yielded peak coordinates at calcarine (peak: -7 -91 -6, T= 14.85) and superior occipital gyrus (peak: -13 -97 19, T= 9.28) regions in the left hemisphere (Table 6).

**Figure 5.**
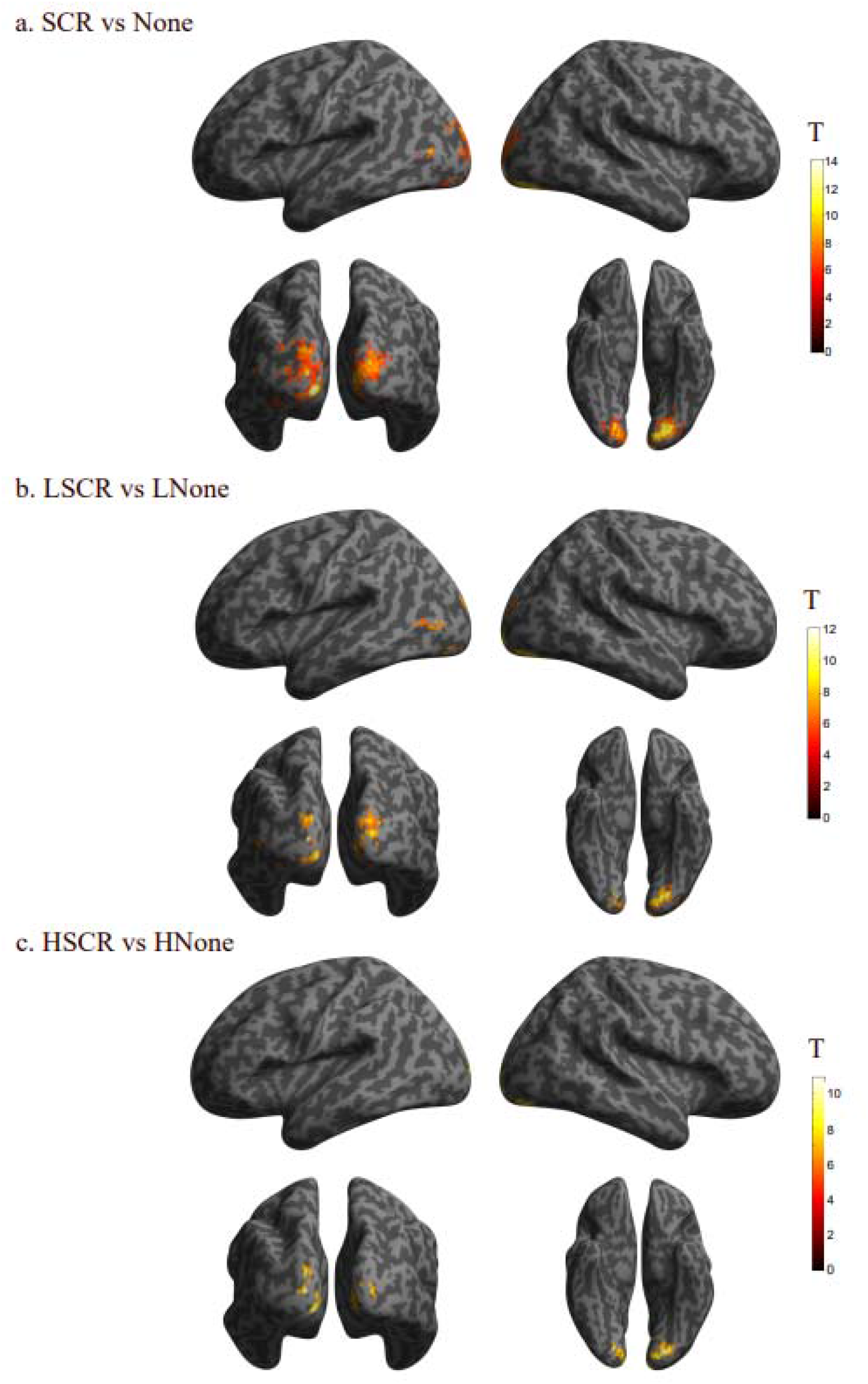
Multivariate pattern analysis results of the attentional load modulation on scrambled BM stimuli. (a) Decoding regions of the SCR versus None classification: Motion sensitive regions in OTC could discriminate SCR and None conditions regardless of the attentional load. (b) Attentional load modulation on SCR was evident: Under low attentional load, SCR and None could be decoded in the motion sensitive OTC regions; while (c) under high load, only the early visual areas could discriminate SCR and None stimuli. Results were FWE-corrected, at a threshold of p <.05.

## 4. Discussion

In this study, the bottom-up processing of biological motion under the top-down modulation of attentional load was measured in order to see whether such socially and ecologically important stimuli as BM would be processed in the BM network even when it was shown away from the focus of attention and if so whether this processing would be modulated by a top-down factor that is attentional load.

In line with the previous literature, our hypothesis on the incidental processing of BM was supported by our findings from the whole-brain univariate activation maps and searchlight based multivariate pattern analysis. Accordingly, BM was processed within the motion-sensitive OTC regions (i.e. hMT+). Moreover, in line with our expectations, this processing was modulated by attentional load as indicated by the decreased activation and diminished decoding regions observed in hMT+. This study has a crucial impact to show the interplay between bottom-up and top-down processing of BM perception through the attentional load paradigm.

### The Bottom-Up Perception of BM

The literature on neural processing of BM under selective attention shows three core brain areas (Thompson & Parasuraman, 2012): (1) The OTC including form and motion sensitive regions (i.e. EBA, FBA, hMT+) as well as STS; (2) parietal (i.e. IPS, IPL); and (3) frontal regions (i.e. IFG, PMC). However, there are inconsistent results in terms of the regions that are associated with the incidental, bottom-up processing of BM (Herrington et al., 2012; Jastorff & Orban, 2009; Saygin & Sereno, 2008).

In the current study, we found that regardless of the attentional load, OTC regions including motion sensitive areas were activated during the bottom-up perception of BM stimuli. These regions were evident specifically under low attentional load. However, under high attentional load, the activation was constrained to the early visual cortex regions. One can make two important inferences based on these findings: (1) In line with the previous literature on incidental processing of BM stimuli, visual feature encoding areas in the OTC were associated with the bottom-up processing of BM; and (2) in addition to the role of OTC in incidental processing, the activation within the OTC was found to be modulated by a top-down factor that is attentional load. Thus, our results suggest that both bottom-up and top-down processing of BM is evident within the OTC. By being the first study to highlight this interaction, this finding emphasizes the need for studying bottom-up and top-down processes together to comprehensivly understand the BM perception more thoroughly.

There were two surprising findings in our results that we did not expect. Firstly, in terms of the recruited brain regions, there was no significant activation found within the pSTS. In the literature, there are inconsistent results towards the neural correlates of bottom-up BM perception. Some studies found activation only in EBA and hMT+ (Herrington et al., 2012; Jastorff & Orban, 2009), while some found additional activity in STS and IPS (Saygin & Sereno, 2008). This disparity between results led us to exploratively hypothesize that under low attentional load, we could see activity maps in not only motion- and form-sensitive areas of OTC, but also in pSTS and parietal cortex. However, our results were constrained to the feature-encoding OTC regions (i.e. hMT+). This finding could be due to the main methodological difference between our study and the study that found pSTS and IPS activity: The stimulus set. In Saygın and Sereno’s retinotopy study (2008), participants’ field of view was populated by intact and scrambled BM stimuli. Meanwhile, in our study and studies of those that did not find pSTS activity (Herrington et al., 2012; Jastorff & Orban, 2009), only one single BM stimulus was shown. Thus, the increased number of stimuli, as well as the contrasting nature of intact and scrambled PLDs might have resulted in stronger stimulation in the BM processing brain regions. Therefore, the possible reason why we did not see pSTS activity in our study could be that using one PLD of BM stimulus may not be strong enough to yield activation beyond visual feature encoding areas in the OTC.

The second surprising finding was that we did not find a significant difference between the intact and scrambled BM stimuli. Based on the literature showing stronger activation towards the intact compared to scrambled BM stimulus within the OTC including STS, we hypothesized that we would find distinct activity maps and stronger activation towards the intact PLDs of BM compared to its scrambled control. However, our results did not yield any difference between scrambled and intact BM stimuli even under low attentional load. This finding may be due to the unsuccessful forming of BM representation indicated by the lack of pSTS activation. The previous literature showed that abstract action representation based on BM stimulus is formed in the pSTS. Moreover, Grossman and colleagues (2004) have found that pSTS activation directly depended on participants’ ability to recognize PLDs as BM. In their perceptual learning study, only when participants could recognize the PLDs of BM stimulus under the noise mask, the pSTS activation was observed. Thus, it could be that our participants could not recognize the PLDs as BM and in turn we neither found difference between intact and scrambled BM stimuli, nor any significant pSTS activation. However, considering the successful completion of participants on a small task conducted prior to the MR scanning in which participants indicated where the intact, not the scrambled, BM was shown on the laptop screen; this conclusion may be unlikely. It would be interesting to replicate our study with a similar stimulus set of Saygın and Sereno (2008) and compare whether pSTS activity and difference between intact and scrambled BM would be observed.

### Engagement of Top-down and Bottom-up Attention Networks

In addition to the bottom-up processing of BM as indicated by the main effect of peripheral stimuli, we have also found a main effect of attentional load evident in distinct network regions. In this study, participants had to direct their attention towards a main visual detection task that was shown at the fovea, while ignoring the task-irrelevant peripheral stimulus. Although the display of the foveal task was kept the same across experimental blocks, the attentional load of the main task was manipulated via increasing the demand of attention resources towards the main task. For the high load task, participants had to detect a conjunction of orientation and color (i.e. upright yellow and upside green t-shapes) rather than a pop-out feature of the t-shape (i.e. color: red shapes). Detecting the conjunction targets rather than the pop-out singleton required participants to focus on the foveal task more than the other (Schwartz et al., 2004). Accordingly, the high load task demanded more attention as compared to low load task. Thus, in line with the previous literature (Schwartz et al., 2005; Desseilles et al., 2009), during the High > Low load contrast (Figure 2.b) as well as the High versus Low load classification (Supplementary Figure 1), we found the fronto-parietal network (FPN) regions indicating increased demand of attentional engagement towards the high load task.

Interestingly, in the Low > High attentional load contrast, we found activation in the default-mode network (DMN) regions. This is the very first evidence to see the interaction between DMN and FPN in an attentional load paradigm. In the working memory literature, it has been shown repeatedly that the DMN activity follows two distinct mechanisms. Firstly, it gets activated during the baseline state in which the participant is not involved in any cognitive task or is engaged in mind-wandering. Secondly, it deactivates during cognitively or attentionally demanding tasks that we see often in the working-memory literature such as performing arithmetics, recalling a set of strings or numbers, or similar to our study, giving response towards a predetermined target (Jenkins, 2019; Arsalidou, 2013; Corbetta, Patel & Schulman, 2008).

When we analyzed the DMN and FPN regions’ activation in High > Low and Low > High contrasts, we found that for the High > Low contrast, there is a significant activation in FPN with a deactivation in DMN regions; and the opposite is correct for the Low > High contrast (Supplementary Figure 2). This finding is in line with the triple network model that was proposed by Mennon (2011), in which as the FPN gets activated DMN regions show deactivation; and vice versa (Lesage & Stein, 2016). Although it was not the aim of this current study to examine the attention networks in depth, one might have expected to see the ventral attention network (VAN, saliency network) rather than the DMN. VAN is the third network in this triple model which is associated with the stimulus-driven processing of the unexpected, task-irrelevant stimuli (Mennon, 2011). The possible reason why we did not observe VAN activity during peripheral stimuli conditions could be that the biological motion stimulus shown at the periphery was not an unexpected event (Corbetta et al., 2000). The amount of trials and the repetition of peripheral stimulus blocks were equal to each other. Also, before the study started, participants were familiarized with the BM stimuli and they were explicitly told that they may see to-be-ignored intact or scrambled BM stimuli at the periphery.

In sum, this is the very first study in the field of attentional load that shows the interplay between DMN and FPN regions. It is not surprising for us to find such a relationship, since in addition to the no-peripheral stimulus baseline condition, we also used a control stimulus (i.e. scrambled motion). This has led us to discriminate the attentional load modulation on the BM stimulus from the distinct brain regions that are associated with low and high attentional load tasks while being able to compare them separately.

### Limitations

In the literature, diminished activation of the task-irrelevant peripheral stimuli has been an indicator of the attentional load modulation. One may suggest that the relatively stronger and extensive activation observed in the low load blocks may be due to the unintentional shift in the eye gaze towards the peripheral stimulus. This possibility could be relevant for our study as well. Considering the early preference and recognition ability of humans towards BM stimuli, one may argue that participants might have shifted their eye gaze towards the peripheral BM stimulus unintentionally. Such shifts in the eye-gaze would yield activity maps similar to the perception of BM under selective attention. However, we did not find any region beyond OTC that could explain the involuntary eye-gaze towards the BM stimulus. So, although we did not use eye-tracker to check whether participants’ eye gaze shifted away from the fixation task, our results show no sign of active viewing of peripheral BM stimuli even under low attentional load.

### Future Directions

Our study shows that the bottom-up processing of BM is supported by motion and form sensitive regions in the OTC. This finding is in line with the model of Giese and Poggio (2003). Accordingly, in their model, there are two separate but parallel processes representing the form and motion pathways. Each pathway consists of an increasing hierarchy of neural feature detectors with increasing receptive field sizes and complexity. The significance of this model is that it clearly supports the immediate recognition, in other words, incidental processing of BM stimuli through feedforward processing. So, observing only the form and motion sensitive areas in the OTC during the bottom-up perception of BM in our study is in line with the model of Giese and Poggio (2003). However, our findings further showed that the OTC regions were modulated by attentional load. This top-down effect of attention on the same brain areas that showed bottom-up processing highlights the possibility of feedback projections. Thus, our findings emphasize the need for more comprehensive models that integrate these two processes and reflect their effect on the overall BM perception. Future studies may look further into building and testing such integrative models.

## 5. Conclusion

In this study we examined the bottom-up perception of BM under attentional load. We have found that even though the selective attention was directed at an attentionally demanding main task at the fovea, the peripheral task-irrelevant BM stimuli were processed incidentally. The neural correlates of this processing yielded activity in motion sensitive regions in the OTC. Thus, our results show that the bottom-up perception of BM is supported by the visual feature encoding areas rather than higher level regions. Finally, the bottom-up processing regions observed in this study were also modulated by attentional load. Thus, our findings add onto the stimulus-driven processing models and studies of BM by showing direct evidence that the top-down and bottom-up perception of BM is interrelated.

## Acknowledgements

H.N. was supported by a graduate student fellowship from a TUBITAK 3501 grant (No: 119K654) awarded to B.A.U. The authors would like to thank Damla Çifçi and Şebnem Türe for their help with data collection, and Murat Batu Tunca and Cem Karakuzu for their help with stimuli design.

