## Supplementary Figures for "Neural processing of bottom up perception of biological motion under attentional load"

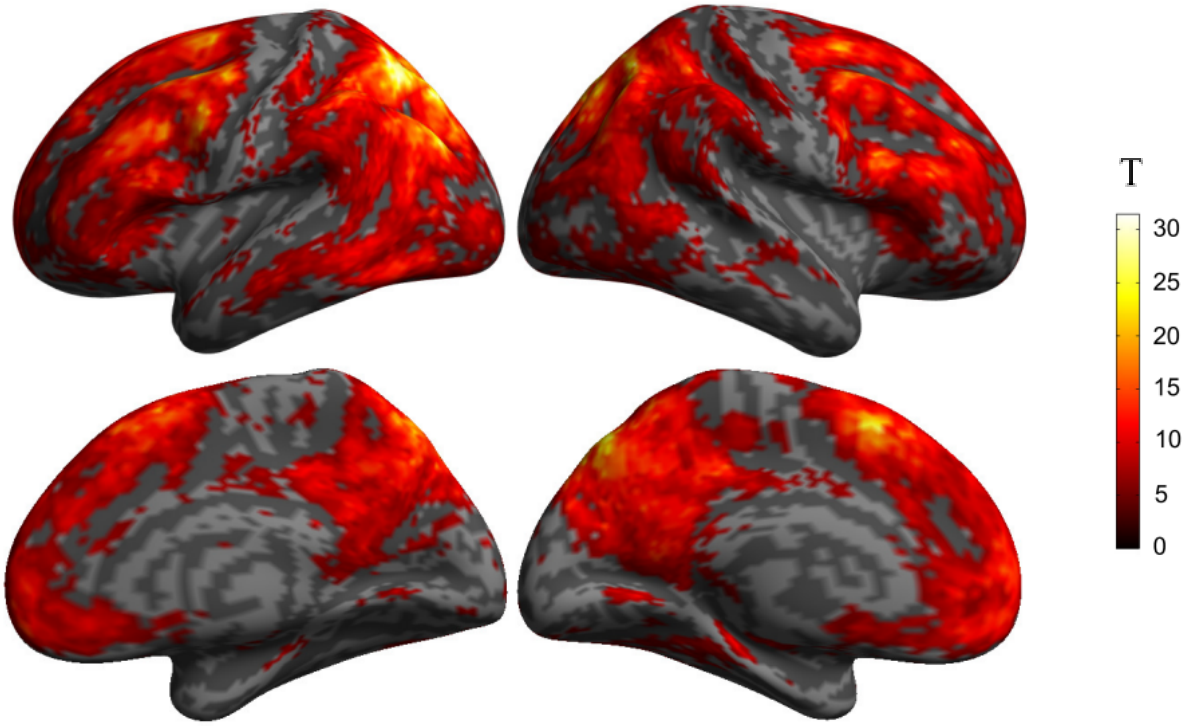


Supplementary Figure 1. High versus Low Attentional Load Decoding Results. MVPA results on high versus low load classification yielded significant decoding results in fronto-parietal network regions.

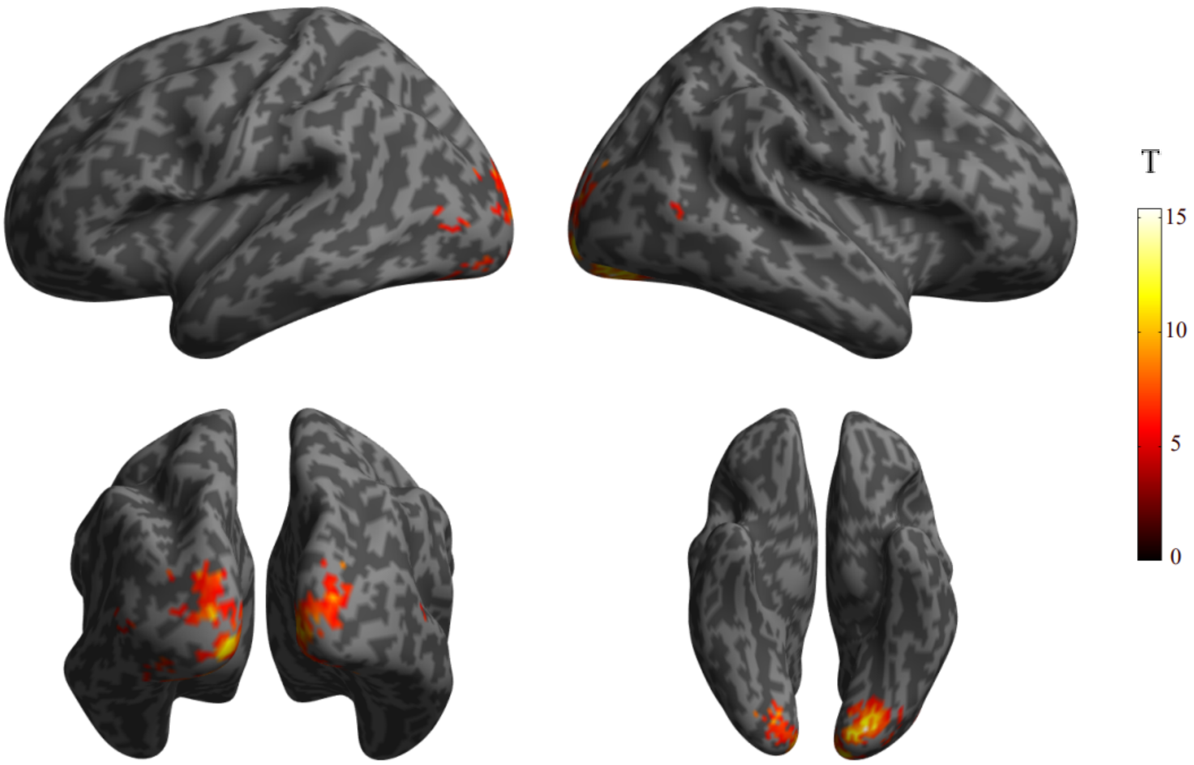


Supplementary Figure 2. Three-way peripheral stimuli decoding maps. MVPA results on threeway BM, SCR, and None classification yielded significant decoding results in OTC regions.

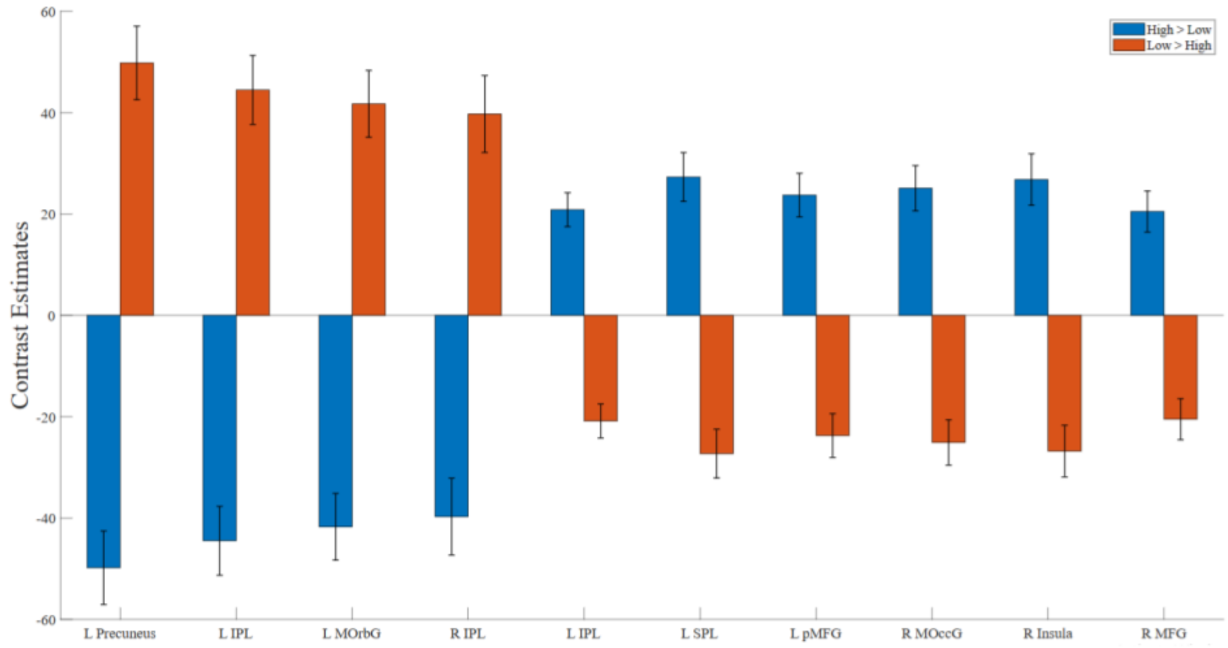


Supplementary Figure 3. DMN and FPN Regions under Low > High and High > Low Contrasts. Under Low > High contrast, FPN regions yielded positive activation while DMN regions were negatively activated. The opposite results were true under Low > High contrast in which DMN regions yielded positive whereas FPN regions resulted in negative activation.
